## Supplementary figures and images for "Loss of full-length dystrophin expression results in major cell-autonomous abnormalities in proliferating myoblasts"

### Supplementary Fig 1

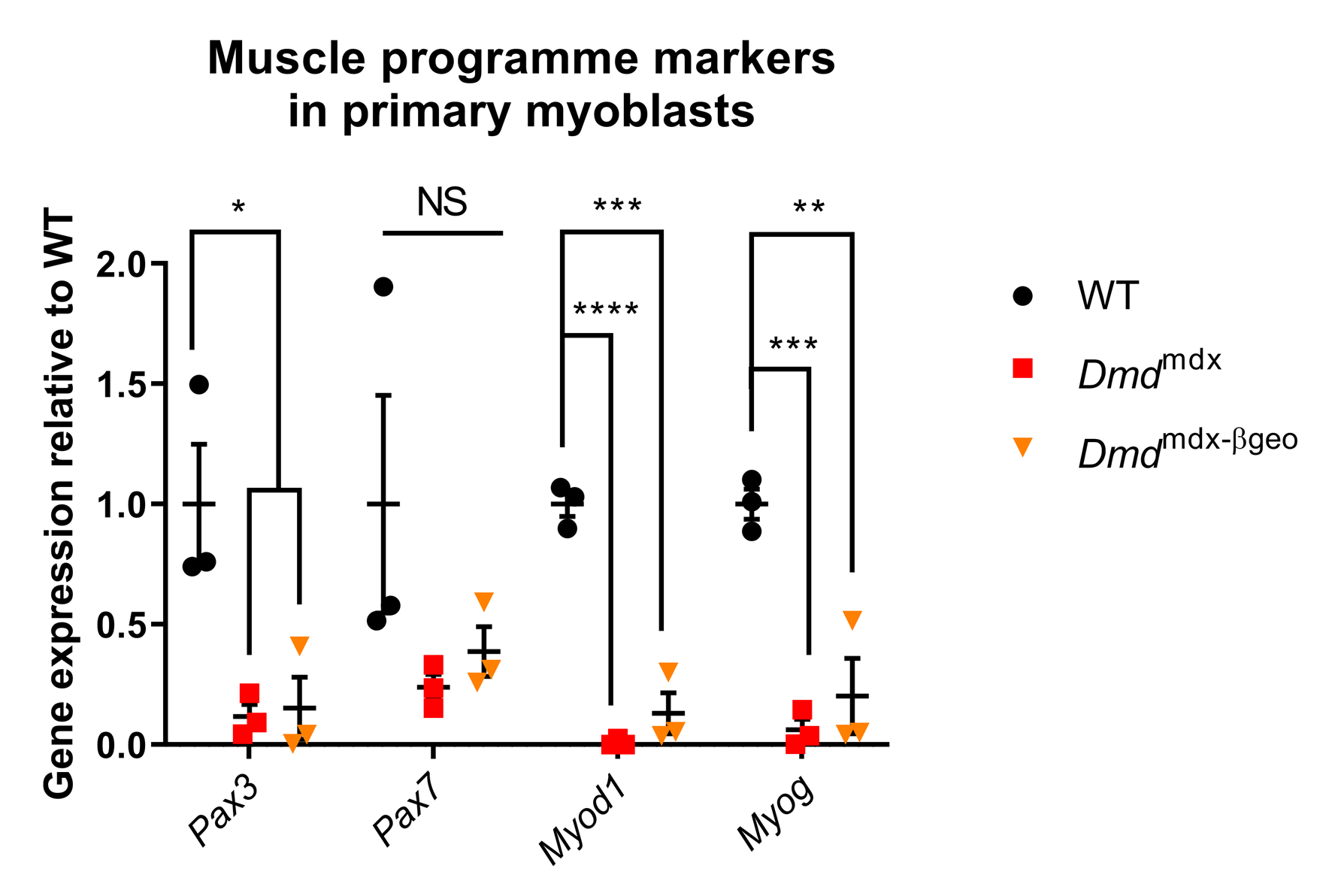

### Supplementary Fig 2

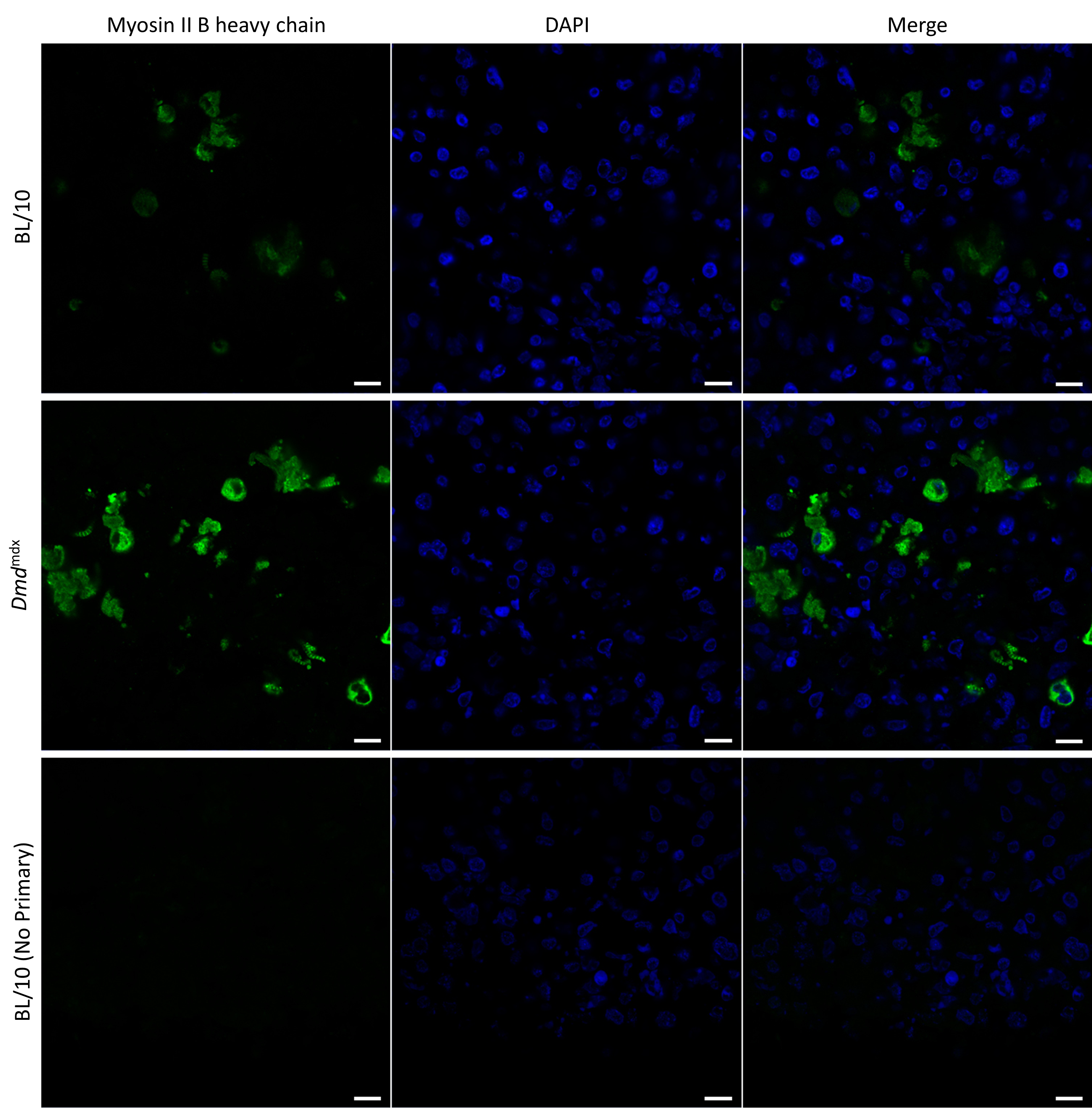

### Supplementary Fig 3

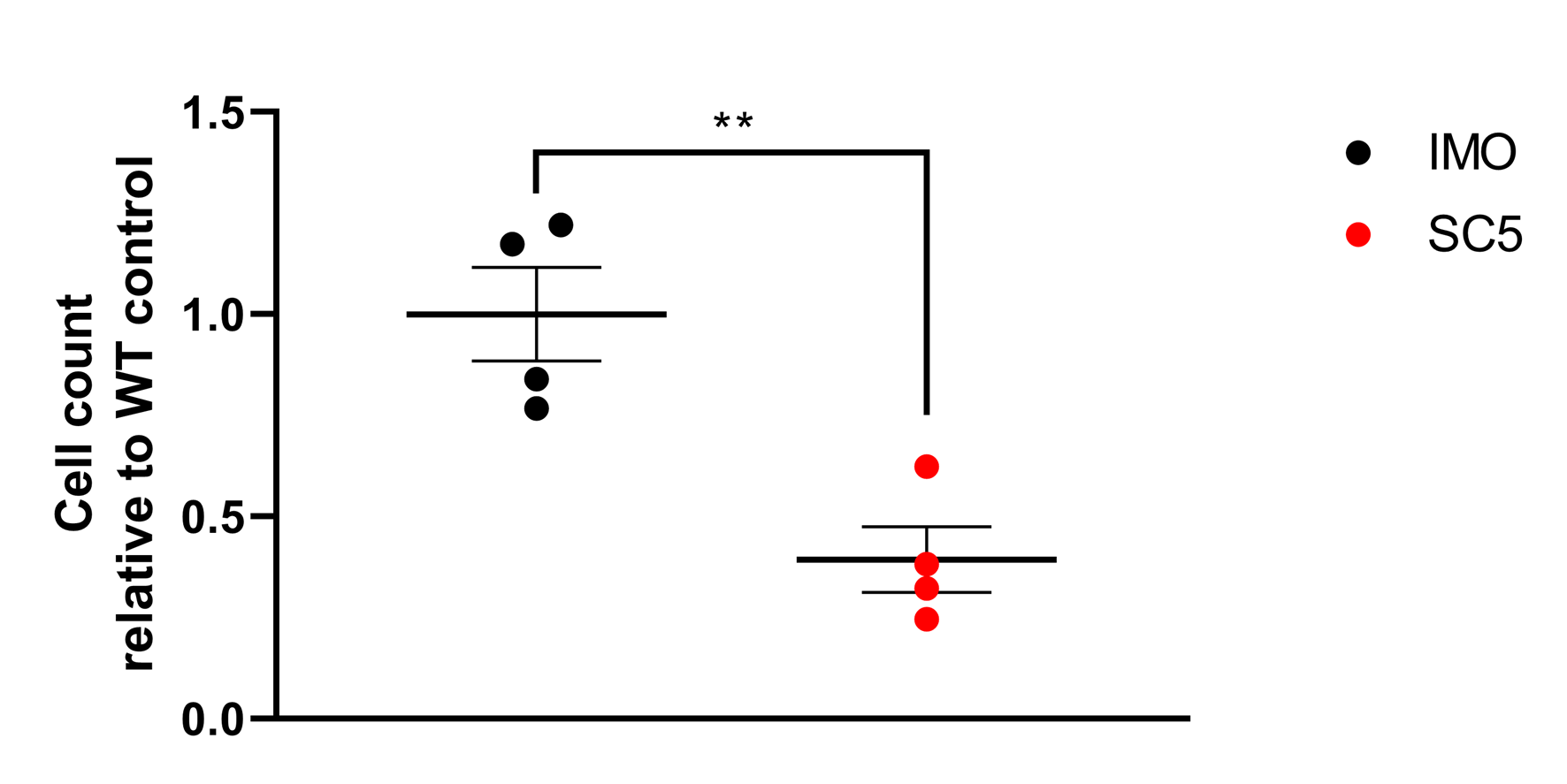

### Supplementary Fig 4

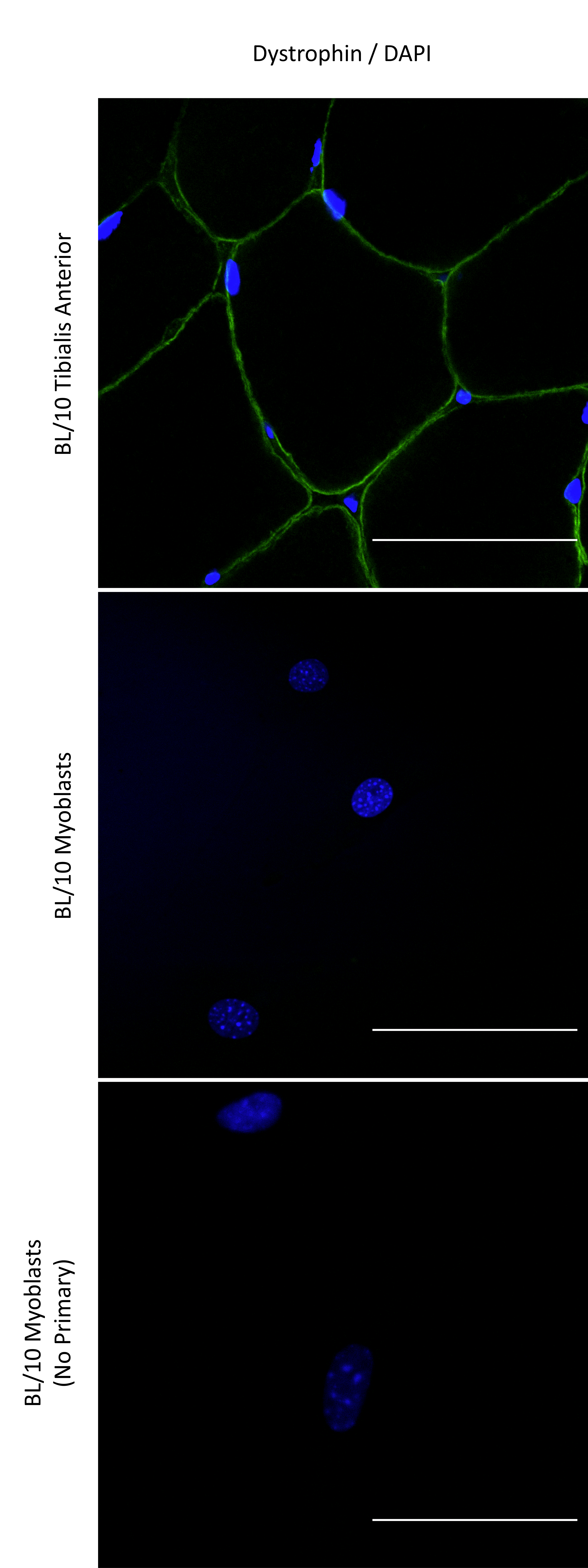

### Supplementary Fig 5

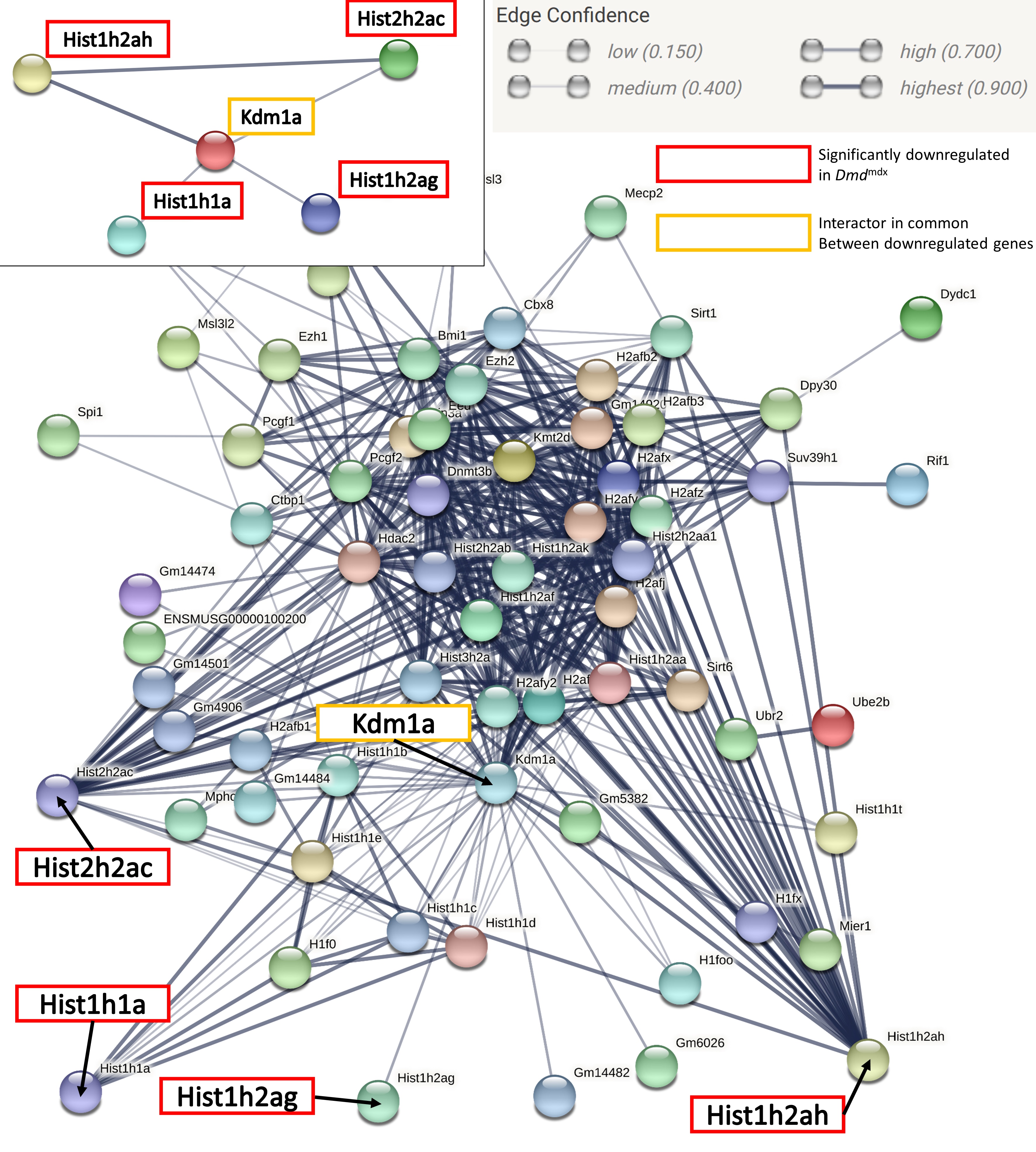

### Supplementary Fig 6

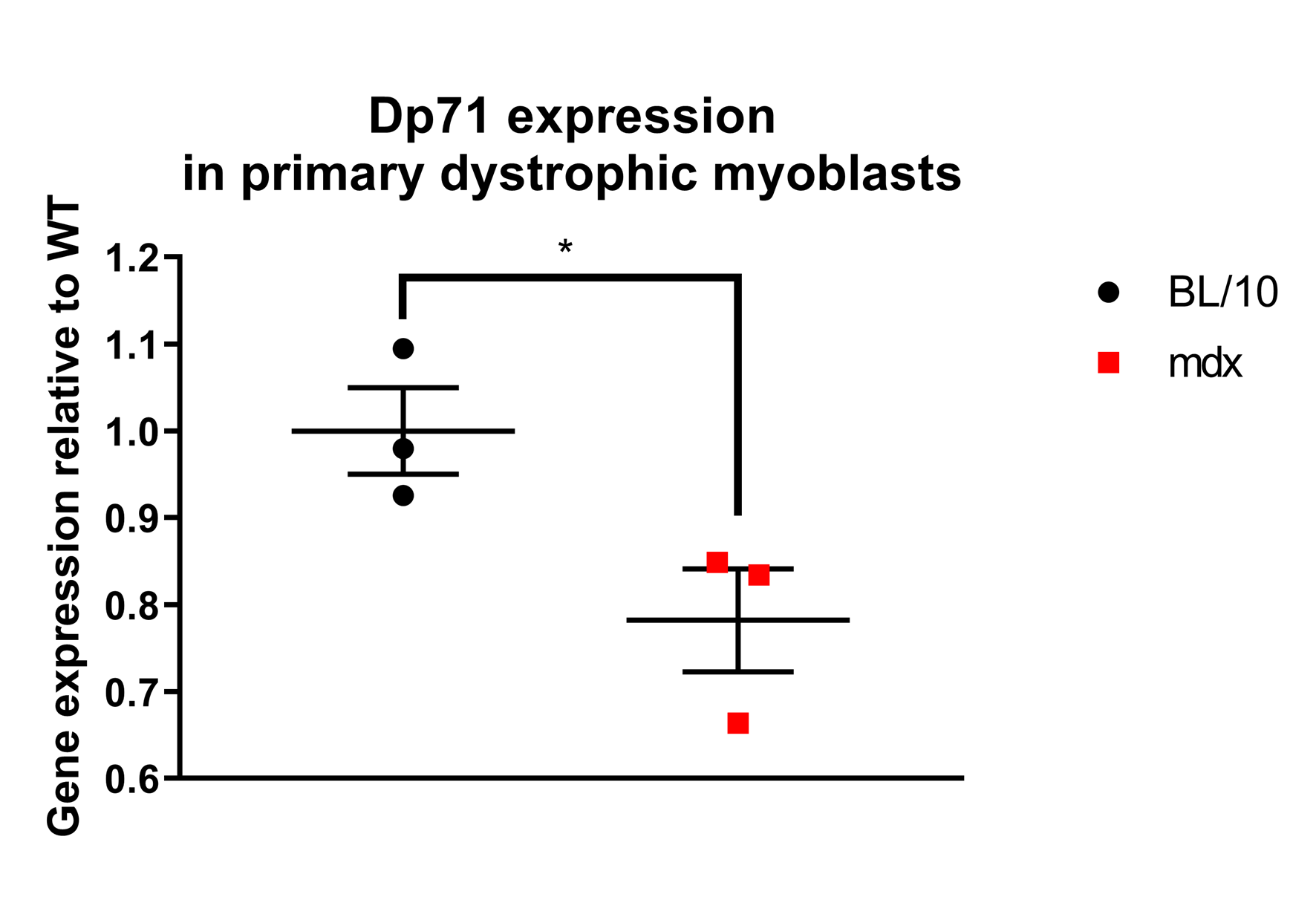

### Supplementary Fig 7

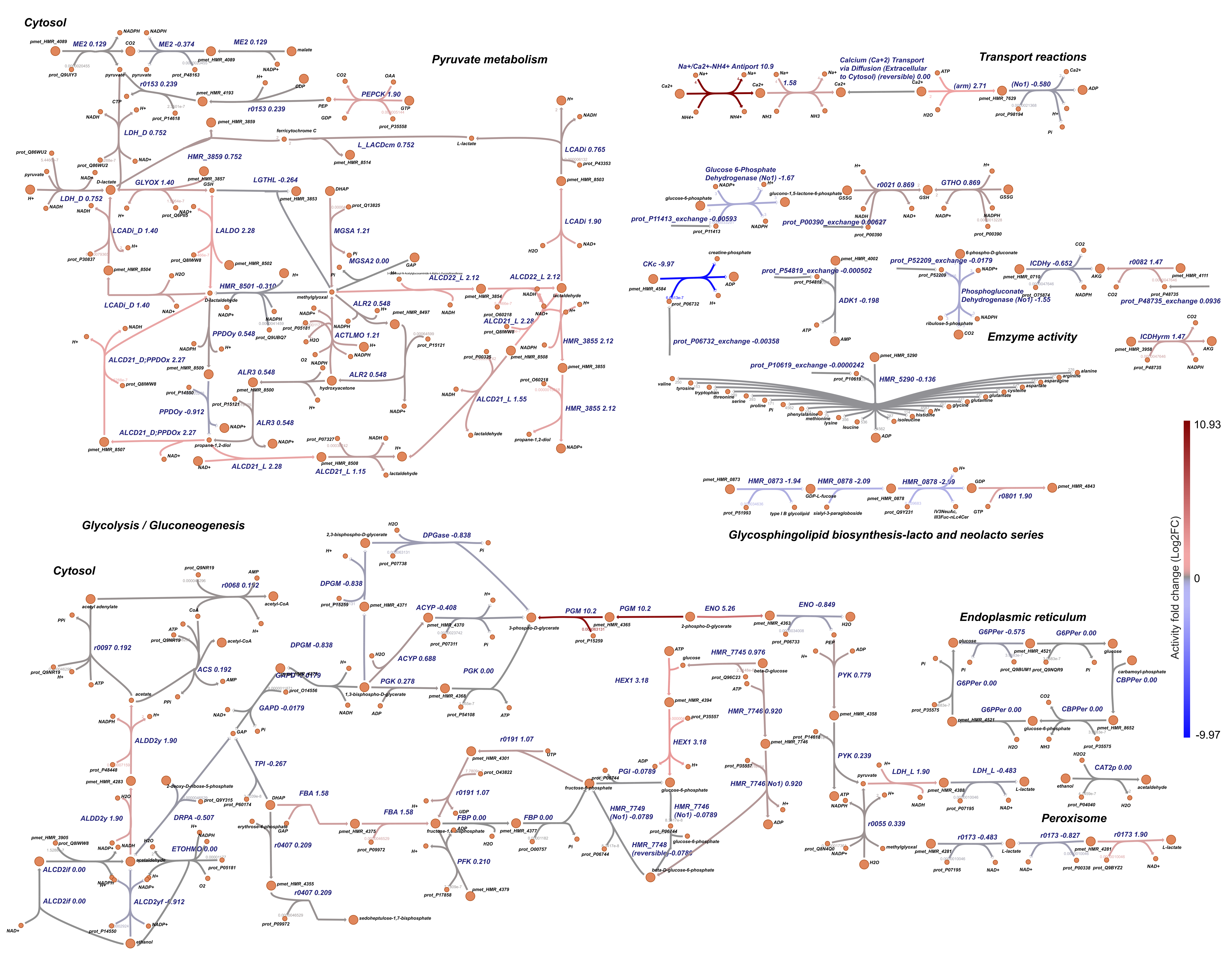

### Supplementary Fig 8

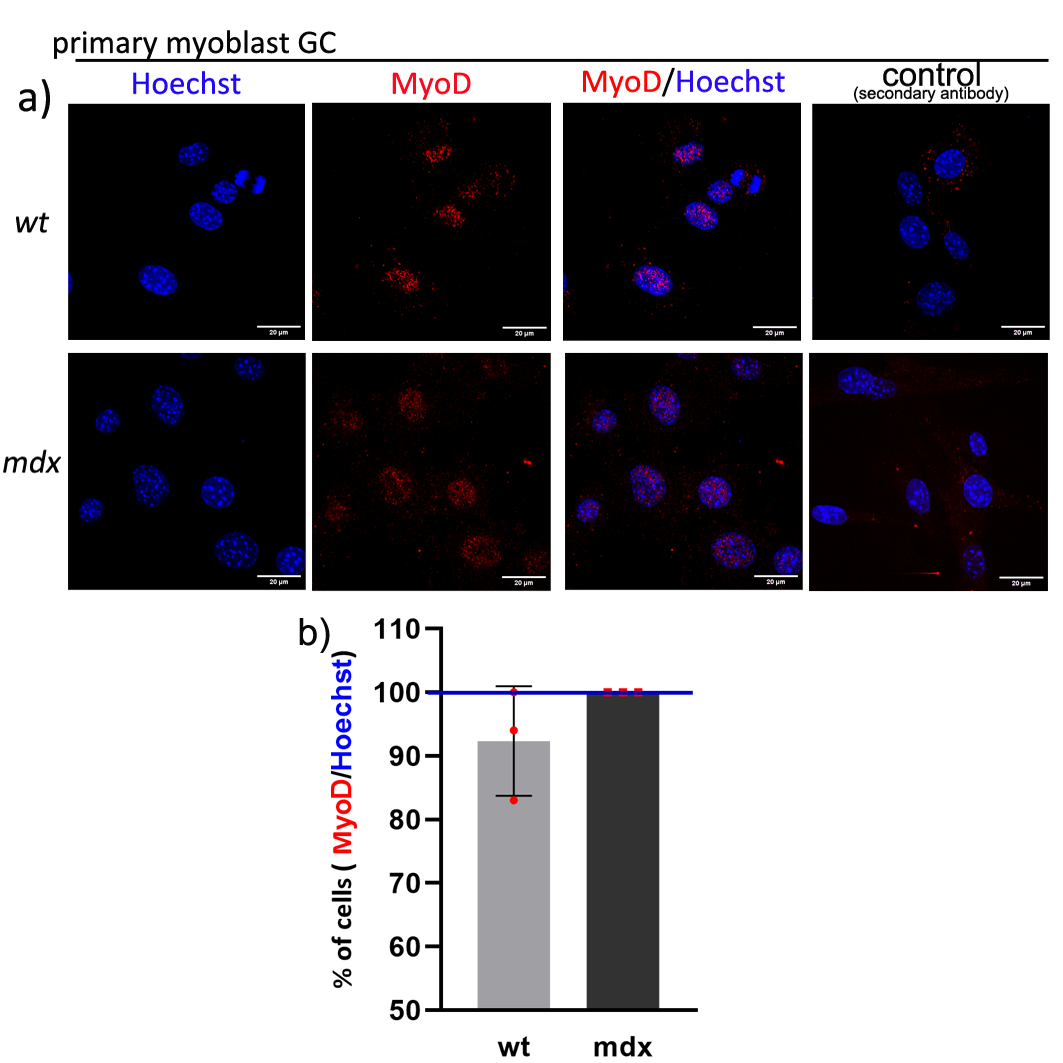
