## Supplementary methods for "Loss of full-length dystrophin expression results in major cell-autonomous abnormalities in proliferating myoblasts"

### Genome-scale metabolic model reconstruction methodology

Significant enzymatic and downstream metabolic changes have been previously observed in DMD^2,3,4,5^. In order to fully take into account both enzymatic and metabolic activity within DMD myoblasts, a condition-specific and enzyme-constrained human metabolic model was constructed to characterise such changes at the genome scale. As a first step, the Human1 genome-scale metabolic model (GSMM) built with GECKO^1,6,7^ was retrieved from (<https://github.com/SysBioChalmers/ecModels/tree/main/ecHumanGEM/model>). GECKO reconstructs the GSMM with enzymatic constraints using kinetic and omic data. This is accomplished by expanding the GSMM's stoichiometric matrix to include rows representing enzymes and columns representing enzyme consumption in reactions, whilst enzyme kinetics (k_cat_ values) are depicted in this matrix by pseudo-stoichiometric coefficients. The enzyme usage in the model is constrained as an upper bound of the respective enzyme (protein) reaction, while the lower bound of those reactions is set to 0. Constraining protein abundance lowers flux variability and improves prediction accuracy^7^.

Flux balance analysis (FBA) is a mathematical method for analysing the flux of reactions within GSMMs. A GSMM is represented as a stoichiometric matrix *S*, with rows corresponding to metabolites and columns representing reactions. At a steady-state, it is assumed that there is no net change in mass/concentration in the system and that mass/concentration is preserved. As a result, the rate of production of each internal metabolite equals the rate of consumption. A column vector *v* reflects the flow through the system (flux rate of each reaction). Under the steady-state assumption, matrix multiplication of the stoichiometric matrix S and column vector v yields the linear equations reflecting the restrictions (*Sv = 0*), as illustrated in Equation 1,

$$\max_{v} c^{T}v$$

$subject to$ $Sv=0$

$$V_{min}\leq v\leq V_{max}$$

(1)

where *S* is a stoichiometric matrix of all known metabolic reactions (metabolites by reactions) and *v* is the vector of reaction flux rates. Additionally, every reaction flux is constrained by lower and upper bounds (*V_min_* and *V_max_*). The linear objective function is represented by the vector c.

Flux Variability Analysis (FVA) is an extended version of FBA, where a range of viable fluxes (i.e. minimum and maximum value) is calculated for each reaction. FVA therefore allows exploring the metabolic range of the cell and does not require an objective function, as it iterates over all the reactions as objectives, calculating their minimum and maximum allowable flux^8^.

Let the vector selecting each biological flux as objective be denoted by *w*. FVA solves two optimization problems (Equation 2) for each flux *v_i_* of interest, after solving Equation 1 with *c = w*.

$\max$/$\min$ $v_{i}$
subject to $Sv= 0$
$w^{T}v\geq\gamma Z_{0}$

$$V_{min}\leq v\leq V_{max}$$

(2)

where $Z_{0}$ = $w^{T}v$ is an optimal solution to Equation 1, and γ is a parameter that controls whether the analysis is performed with respect to suboptimal network states (*0 ≤ γ ≤ 1*)^9^.

The fold change value of RNA-Seq data obtained using DeSeq2 (as described in Section RNA Sequencing), was used to further constrain the GSMM. Using a previous method for building context-specific models^10^, the RNA-Seq data was integrated into the GSMM to obtain a myoblast-specific metabolic network^8^. The idea is to constrain the base enzyme constrained human metabolic model with upper and lower bounds on metabolic reactions depending on the fold change value of gene expression data, as illustrated in Equation 3.

$$lb\left( i \right)=lb\left( i \right)*\left( rxnExprFC\left( i \right)^{\theta} \right)$$

$ub\left( i \right)=ub\left( i \right)*\left( rxnExprFC\left( i \right)^{\theta} \right)$

(3)

where *lb* is the lower bound and *ub* is the upper bound of the reaction, *i* is the index of the reaction, *rxnExprFC* represents a fold change of each reaction, calculated using the fold change values of the gene expression and taking into account the role of each gene in each reaction^11,12^. *θ* is the hyperparameter used to constrain the model based on the gene expression fold change value. Here, *θ* was set to 2.

In addition to the transcriptional constraints described above, we implemented additional constraints based on the literature. Significant enzymatic and metabolic disruption has been shown in DMD, where modest enzymatic alternations may be observed in the early stages but become more extensive with the progression of muscle tissue degeneration^13,14^. A significant upregulation in glycolytic enzymes, including Hexokinase-1 and Pyruvate Kinase M2, was observed in dysfunctional muscles, indicating the increased glycolytic activity^15^. The activity of individual glycolytic enzymes (such as alpha-glucan phosphorylase, phosphoglucomutase, and aldolase), creatine kinase and Adenylate kinase 1 is reduced in DMD^5^. Furthermore, the first two enzymes of the pentose phosphate pathway of glucose utilisation, glucose-6-phosphate dehydrogenase and 6-phosphogluconate dehydrogenase, have enhanced activity. The activities of lysosomal cathepsin enzymes (such as cathepsins D, A, B1, C, and dipeptidyl peptidase II; protein hydrolyzing enzymes) are also enhanced in muscular dystrophies. Based on these increased or decreased enzymatic activities from the available literature on metabolic alterations, the upper and lower bound of the specific metabolic reactions associated with these enzymes were further constrained as follows.

Firstly, FBA was computed for all reactions using biomass as an objective function. Then the flux rate calculated for the reactions associated with these enzymes was used as a value for the upper bound to ensure reduced enzymatic activity, or as the lower bound to ensure enhanced enzymatic activity, following the cases above. Furthermore, we constrained the biomass output to at least 50% of its maximum growth.
